# Putative interfollicular stem cells of skin epidermis possess a specific mechanical signature that evolves during aging

**DOI:** 10.1101/2025.02.12.637845

**Authors:** Sarah Miny, Gaël Runel, Julien Chlasta, Jean-André Lapart, Christelle Bonod

**Affiliations:** Skin Functional Integrity group, Laboratory for Tissue Biology and Therapeutics Engineering (LBTI), CNRS UMR5305, University of Lyon, France; BioMeca, Lyon, France

**Keywords:** Skin, stem cells, aging, dermo-epidermal junction, biomechanical properties, AFM

## Abstract

Skin homeostasis and self-renewal are partially maintained by interfollicular stem cells (ISCs), located in the basal layer above the dermal papillae of the dermo-epidermal junction (DEJ). Aging leads to a decline in skin renewal and a concurrent reduction in stem cell potential. It is also marked by disorganization of the extracellular matrix in both the DEJ and dermis, and flattening of the DEJ. To better understand ISC aging, new methods are needed to characterize ISCs and their environment. Since mechanical properties of cells and their substrate influence cell fate, we employed atomic force microscopy to explore whether ICSs niches and the DEJ exhibit distinct mechanical properties. Our findings reveal that ISCs possess greater stiffness than other basal cells, a mechanical signature that diminishes with age. Additionally, the DEJ beneath ISCs shows higher stiffness than under other basal cells, providing ISCs with a specific mechanical environment, which also deteriorates during aging. *In vitro*, sorting of ISCs based on MCSP expression effectively isolates ISCs beneath the dermal papillae, allowing the measurement of their mechanical signature and stemness potential under varying mechanical conditions. The study of ISC mechanical signatures offers a promising approach for characterizing 3D skin models and understanding defects in skin renewal and wound healing.

## INTRODUCTION

The skin, the largest organ of the human body, serves as the primary barrier against environmental factors. The epidermis, its outermost layer, possesses a remarkable ability to self-renew, which is critical for skin homeostasis and for repairing wounds. This regenerative function is supported by epidermal stem cells located in the basal layer of the epidermis [1]. Stem cells play a key role in maintaining tissue homeostasis. In humans, the basal layer, situated above the dermo-epidermal junction (DEJ), harbors undifferentiated keratinocytes, transit amplifying (TA) cells, and a population of putative interfollicular stem cells (ISCs) [2], [3]. The DEJ, composed of specialized extracellular matrix (ECM) proteins, such as collagen VII and XVII or laminin 332 [4], [5], anchors epidermal cells to the dermis and facilitates communication between the dermis and basal epidermal cells [6][7], transmitting signals essential for proliferation and differentiation. The DEJ is characterized by its undulating topography, with structures known as dermal papillae (DP) and epidermal rete ridges (RR). ISCs, situated specifically above the DP, express key stem cell markers, including Melanoma Chondroitin Sulfate Proteoglycan (MCSP), Leucine-rich Repeats and Immunoglobulin-like domain protein 1 (LRIG1), and elevated levels of β1-integrin [8], [9], [10], [11]. Single-cell RNA sequencing has revealed four spatially distinct stem cell populations within the basal layer, residing at the top and bottom of the rete ridges, underscoring the heterogeneity of human interfollicular epidermal stem cells [12]. Several studies have demonstrated that stem cells can be isolated from the basal layer through flow cytometry, by selecting cells with high α6-integrin (CD49) expression and low transferrin receptor (CD71) levels. These cells exhibit significant clonogenic potential and long-term self-renewal in culture [13], [14], [15]. Similarly, basal cells expressing high levels of β1-integrin and MCSP demonstrate robust self-renewal and clonogenic capabilities *in vitro*. Cluster of stem cells marked by MCSP are thought to enhance cell-to-cell adhesion, contributing to the organization of these cells [16]. The combined expression of α6-integrin, CD71, and MCSP presents a promising method for isolating interfollicular stem cells localized specifically above dermal papillae.

Stem cells typically reside in specialized microenvironments, known as niches, which play a pivotal role in regulating their behavior and fate [17]. In the case of ISCs, their interaction with proteins of the DEJ, where ISCs are anchored, contributes to the modulation of their niche [18] [19]. Laminin 332, for instance, interacts strongly with integrin on the surface of ISCs, helping to maintain epidermal homeostasis by influencing the balance between stem cell renewal and differentiation [20], [21]. The properties of the cellular substrate-its composition, stiffness, and topography-are also integral to the niche environment, influencing stem cell fate and renewal capacity. For example, mesenchymal stem cells tend to differentiate on softer substrates [22], while dental pulp stem cells show reduced proliferation under similar conditions [23]. Conversely, keratinocytes and glioma cells grown on rigid substrates demonstrate greater proliferative potential [24], [25]. Keratinocytes also adapt to changes in substrate stiffness by forming robust, interconnected networks of keratin filaments, thereby increasing their cellular rigidity [26]. The rigidity of the cell’s cytoskeleton has been shown to correlate with specific differentiation states, serving as an indicator of cell fate [27], [28].

The attachment of ISCs to the DEJ is linked to cytoskeletal architecture and cell morphology, which are crucial for maintaining their stemness, as well as regulating their proliferation and differentiation [7], [18]. During aging, the skin undergoes numerous structural and functional changes. The DEJ flattens, and its protein composition is altered, with decreased expression of laminin 332, β4-integrin, and collagen types IV, VII and XVII [29], [30], [31], [32]. These changes contribute to a decline in the rate of epidermal renewal, impaired wound healing, and a thinning of the epidermis [33], [34]. Additionally, the stemness potential of ISCs diminishes with age, as evidenced by reduced expression of β1-integrin and MCSP in the basal layer cells [35] [8].

There is a growing need for new methods to characterize ISCs and their environment during skin aging in order to understand the defects associated with this process. Many genetic skin diseases and skin aging are linked to altered cellular or tissue mechanical properties of the skin [36], [37], [38]. Atomic Force Microscopy (AFM) is one of the current methods used to study the mechanical properties of tissues, cells, or subcellular structures at nanometer resolution [39], [40], [41], [42][41][39]. To date, the mechanical characterization of ISCs within tissue has not been studied. Understanding the mechanical properties of ISCs would provide a novel approach to characterizing ISCs and their environment. The aim of our study was to investigate whether ISCs, and their substrate, the DEJ, have a specific mechanical signature and whether this evolves during aging. We then sought to specifically isolate ISCs located above dermal papillae from human skin, to characterize them functionally and mechanically *in vitro*. Our results showed that ISCs and their environment have a distinct mechanical signature, which is lost during skin aging, thus highlighting a potential skin aging model based on the evolution of the mechanical signature of ISCs.

## METHODS

### Human skin samples

Human skin tissue explants were collected with informed consent from patients undergoing surgical procedures, following the ethical guidelines of Biopredic International. The donors were healthy Caucasian females, Phototype III or IV, aged between 20 and 78 years. After collection, the hypodermis was carefully removed using scissors and a scalpel. The skin was used for keratinocyte extraction or cut into 1 cm-diameter round pieces, flash-frozen in liquid nitrogen, and stored at −80°C. Frozen samples were embedded in OCT embedding matrix (Sakura 4583), and 16 µm cryosections were prepared using a Cryostat (LEICA CM3050S) at −20°C. The sections were mounted on SuperFrost Plus slides (Fisher) and stored at −20°C.

### Immunofluorescence assays

Cryosections were thawed and rehydrated in 1X PBS for 10 minutes. Sections were then fixed in 4% formaldehyde (Sigma) diluted in 1X PBS for 20 minutes. Following three washes with 1X PBS, sections were blocked with 5% BSA (Sigma) in 1X PBS for 2 hours at room temperature (RT). Primary antibodies, diluted in 5% Bovine Serum Albumin (BSA), were applied overnight (O/N) at 4°C. The primary antibodies used included mouse anti-MCSP (NG2) at a 1:100 dilution (Santa cruz_SCs-80003) and mouse anti-collagen VII (Santa cruz_SCs-33710). After three washes with 1X PBS, sections were incubated with secondary antibodies for 2 hours at RT. Secondary antibodies, diluted in 5% BSA, included anti-goat CF555 (1:800, Sigma_Sab4600072), anti-mouse Alexa Fluor488 (1:800, Thermofisher_A28175), anti-mouse IgG2a Alexa Fluor555 (1:200, Invitrogen A-21137), and anti-mouse IgG1 Alexa Fluor488 (1:200, Invitrogen A-21121). Nuclei were stained with DAPI (1 µg/mL, Sigma) for 20 minutes at RT. Finally, sections were washed three times with 1X PBS and stored in 1X PBS at 4°C. Mounting was performed using a 1X PBS-50% glycerol solution (Euromedex) as the mounting medium between the slide and coverslip, which were sealed with nail varnish and stored at 4°C.

### Image acquisition and analysis

Fluorescence imaging was conducted using a ZEISS LSM880 confocal microscope (CIQLE-Centre d’Imagerie Quantitative Lyon Est, Lyon 8) equipped with a 40× 1.3 Oil Plan-Apochromat objective. Image analysis was performed using ImageJ software.

### Atomic Force Microscopy (AFM) measurements

AFM measurements were conducted using a Resolve Bioscope (Bruker Nano Surface, Santa Barbara, CA) mounted on an inverted optical DMI8 microscope (Leica). A conical attached to a flexible cantilever with a spring constant of 0.35 N/m (DNP-10A, Bruker AFM probes) was utilized for the measurements. Prior to each experiment, the deflection sensitivity of the cantilever was calibrated, and the spring constant was determined using the thermal tuning method. Data acquisition was performed with Nanoscope software (version 9.1.R.3) in AFM QNM (Quantitative Nanomechanical Mapping) mode. Force measurements were conducted in 1x PBS on human cryosections that had been fixed with 4% formaldehyde and immunolabeled for MCSP and collagen VII. Force volume acquisitions were carried out over a 25×25 μm area at three distinct locations: (1) at the level of MCSP-positive cells and putative ISCs above the DP; (2) at the level of MCSP-negative cells above the RR; and (3) in the reticular dermis. This 25×25 µm scan size allowed for the characterization of the mechanical properties of both basal epidermal cells and the ECM composing the DEJ and papillary dermis (Figure 2). Each force volume included 4,096 measurement points, with each point corresponding to an indentation force curve from which the elastic modulus (Ea) was extracted. The quantification of the elastic modulus from the raw force curves was performed using the Sneddon mathematical model within BioMeca Analysis processing software. Finally, stiffness maps were reconstructed from the elastic modulus values, allowing for the extraction of the stiffness (Figure 2).

### Keratinocyte extraction and culture

Primary keratinocyte cultures were obtained with informed consent from patients undergoing surgical discard, adhering to the ethical guidelines of the Cell and Tissue Bank of Hospices Civils de Lyon. The donors were healthy Caucasian females aged between 20 and 78 years, all classified as phototype III. Following the manual removal of the hypodermis, skin explants were cut into 0.5 cm² pieces and incubated overnight at 4°C in a solution containing 20 U/ml Dispase II (Roche Diagnostics), 10,000 U/ml penicillin, and 10,000 mg/mL streptomycin (P/S; Gibco), 0.02 mg/ml white trypsin (Gibco), and Dubelcco’s Modified Eagle’s Medium (DMEM; Gibco). After incubation, the epidermis was separated from the dermis and incubated for 20 minutes a at 37°C in 1X EDTA trypsin solution (Trypsin-Ethylene Diamine Tetraacetic Acid; Gibco) to dissociate epidermal cells. The dissociated cells were then cultured in KBM Gold (Keratinocyte Growth Medium BulletKit, Lonza) and counted using a Malassez chamber with trypan blue (Sigma Aldrich) for viability assessment. Following cell sorting, the keratinocytes were seeded on rat collagen I matrix (100 µg/mL) at a concentration of 40.000 cells/cm² in KBM Gold medium. The cultures were incubated at 37°C in a 5% CO2 atmosphere. The medium was changed the following day and refreshed every two days thereafter.

### Flow cytometric isolation of interfollicular stem cells situated above the dermal papillae

Freshly isolated keratinocytes were stained with a panel of antibodies in 100 µl of FACS buffer. Initially, the cells were incubated for 30 minutes at 4°C with the primary antibodies mouse anti-MCSP (SCs-80003, Santa Cruz). Following this, the cells were washed and incubated for an additional 30 minutes at 4°C with the secondary antibodies, anti-mouse IgG2a Alexa Fluor555 (A-21137, Invitrogen). After another washing step, the cells were stained for 30 minutes at 4°C with R-Phycoerythrin (R-PE)-conjugated rat anti-human CD49f antibodies (561894, BD Biosciences) and APC-conjugated mouse anti-human CD71 antibodies (17-0719-42, Invitrogen). Following this staining procedure, the cells were washed again and stored in FACS buffer at 4°C until sorting. Just prior to sorting, the cells were labeled with 1X DAPI (Sigma) to identify the nuclei. Using a BD FACS ARIA II cell sorter (SFR BioSciences, France), the following populations were isolated: CD49high-CD71low-MCSPhigh (MCSP+ SCs), CD49high-CD71low-MCSPlow (MCSP-SCs), and CDhigh-CD71high-MCSPhigh (MCSP+ TAs) (Figure supp 1). After sorting, the cell populations were counted, and their viability was assessed using trypan blue.

### Statistical analysis

Data are presented as mean ± standard deviation (SD). Statistical significance was determined based on the normality of the data, assessed using the Shapiro-Wilk test. For normally distributed data, either the Student t-test, one-way or two-way analysis of variance (ANOVA) was employed using Prism software (version 10, GraphPad Software, San Diego, CA, USA). In cases where the data did not meet normality assumptions, the Mann-Whitney or Wilcoxon test was applied. Differences between means were deemed statistically significant at p < 0.05, with thresholds of *p < 0.05 and **p < 0.01 indicating varying levels of significance.

## RESULTS

### Immunofluorescence-based characterization of putative ISCs

To identify ISCs in the basal layer of human skin cryosections, we employed an immunofluorescence approach, specifically targeting the MSCP protein. Additionally, collagen VII labeling was used to visualize the DP and RR, while nuclei were stained with DAPI. In sections from a 36-year-old donor, MCSP labeling revealed a distinct localization of ISCs in clusters of basal cells situated above the DP (Figure 1A). This consistent localization of MCSP was observed across all samples, confirming the expected positioning of ISCs above the DP.

**Figure 1.**
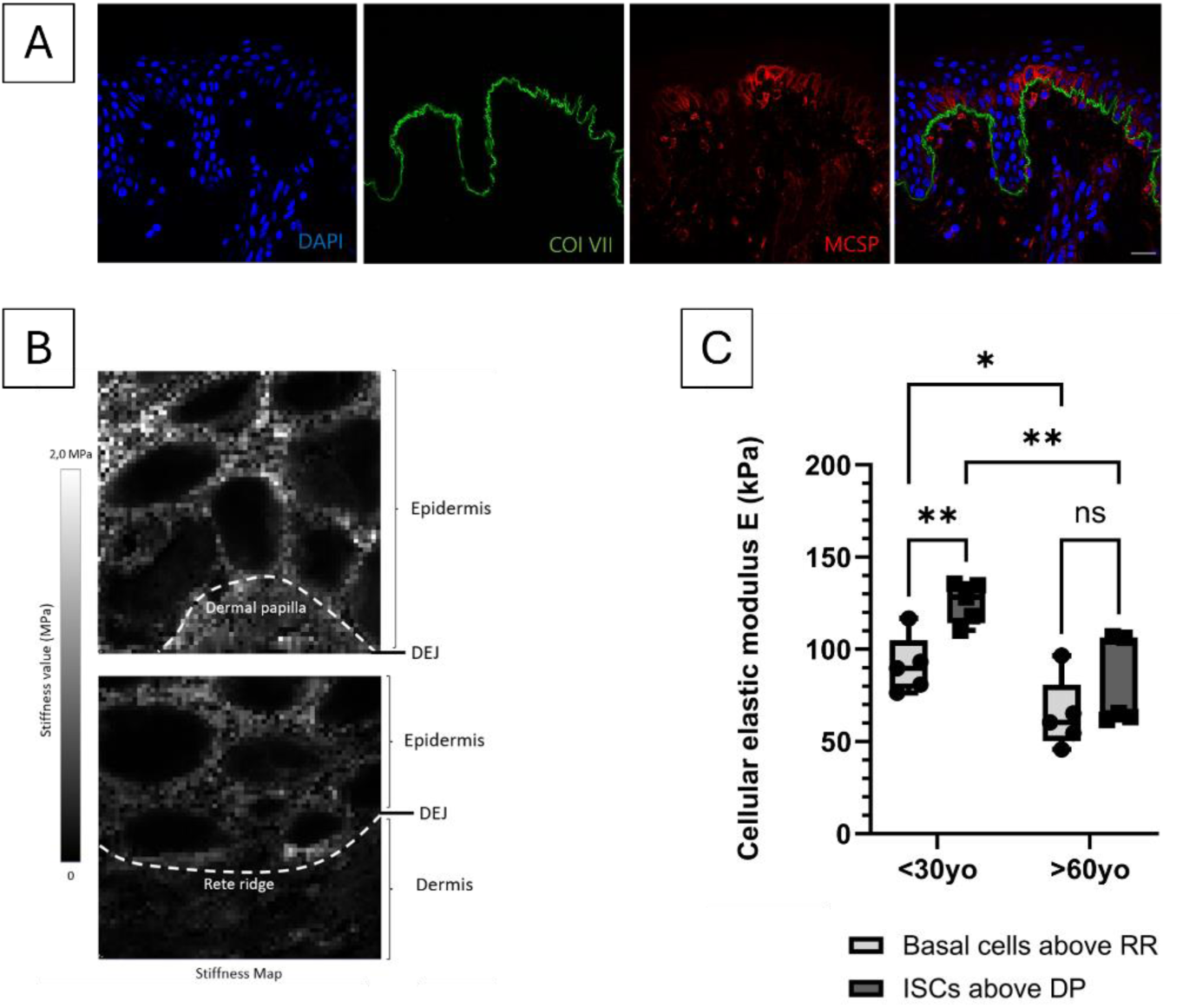
Analysis of average cell stiffness measured by AFM on frozen epidermal sections from human skin of different ages. The study included a group of donors under 30 years old (N=5; ages: 22, 24, 25, 27, 29) and a group over 60 years old (N=5; ages: 60, 60, 61, 62, 64), with three sections analyzed for each donor and more than 15 AFM measurements taken per donor. For the AFM measurements, three independent areas were assessed per section for each donor, averaging three AFM measurements of 25 x 25 µm at the level of ISCs and at the level of cells above the RR. **(A)** Representative tissue sections the group under 30 years stained for collagen VII (green), MCSP (red), and DAPI (blue). Scale bar = 20 µm. **(B)** Stiffness maps reconstructed from the elastic modulus values extracted from indentation force curves on human sections at the level of the DP and RR. Scale bar = 5 µm. **(C)** Box plot illustrating the average cell stiffness of ISCs above the DP (dark grey) and basal layer cells above the RR (light grey) for both age groups: under 30 years (N=5) and over 60 years (N=5). A two-way ANOVA with Tukey’s multiple comparison test was employed to compare cellular stiffness between ISCs and basal cells above the RR in young and older human epidermal sections. ns = not significant; * p-value < 0.05; ** p-value < 0.01.

### Mechanical properties of ISCs

To investigate whether putative ISCs exhibit distinct mechanical properties, we employed an AFM-based approach to assess cell stiffness. This method enables the detection of specific cellular stiffness across different subpopulations within the basal layer. Our AFM measurements focused on putative ISCs, identified by their expression of MCSP, located above the DP, as well as on the MCSP-negative basal cells situated above the RR. This comparative analysis allowed us to generate stiffness maps, from which we extracted the stiffness values of the various basal cell populations (Figure 1B). In samples from individuals under 30 years of age, we observed a significant difference in cell stiffness among the different populations. Specifically, the average cell stiffness of ISCs above the DP was approximately 1.4 times higher than that of the basal layer cells above the RR (Figure 1C). These findings indicate the presence of specific mechanical properties that distinguish these cell subpopulations in younger skin, highlighting that the stiffness of putative ISCs is greater than that of the basal layer cells located above the RR.

### Impact of skin aging on the mechanical properties of putative ISCs

During skin aging, a decline in stem cell potential has been reported, correlating with a reduction in stem cell markers such as β1-integrin and MCSP [8]. This prompted our investigation into whether ISCs exhibit distinct mechanical signatures during aging.

For that, AFM measurements were performed on samples from individuals 60 years old, specifically targeting the putative ISCs located above the DP and the basal cells above the RR. In the samples from this age group (Figure 1C), no significant difference in cell stiffness was observed between the two populations. The average stiffness of ISCs above the DP was comparable to that of basal layer cells above the RR. Next, we compared the cellular stiffness of ISCs in skin samples under 30 years to those over 60 years. The results revealed that the average cellular stiffness of ISCs in the younger group was significantly higher than that of ISCs in the older group (Figure 1C). This suggests a notable modification in the mechanical signature of ISCs with aging. A similar trend was observed in the stiffness of basal cells above the RR. In conclusion, our findings indicate that aging is associated with a significant decrease in the cellular stiffness of ISCs, leading to a loss of their distinct mechanical signature, as they exhibited mechanical properties comparable to those of basal cells above the RR.

### The stiffness of ISCs is linked to the topography of the DEJ

Previous studies have shown that the differentiation of epidermal stem cells can be influenced by the specific topography of their substrate [43]. Therefore, we aimed to investigate whether the topography of the DEJ correlates with the stiffness of ISCs. To characterize the DEJ topography, we measured the height of the DP in skin sections from individuals under 30 years old and over 60 years old (Figure 2A). Collagen VII labeling was employed to visualize the DEJ. Our findings revealed that the average height of the DP in skin samples from individuals under 30 years is significantly greater than that in samples from those over 60 years (Figure 2B), confirming the flattening of the DEJ with age. Next, we analyzed the correlation between DP height and ISCs stiffness for each donor. The Pearson correlation coefficient (r = 0.6583) indicated a significant positive correlation between DP height and ISCs stiffness (Figure 2C). In contrast, no such correlation was found between DP height and the stiffness of basal cells above the RR. Thus, we established a clear correlation between DEJ topography and the mechanical properties of ISCs.

**Figure 2.**
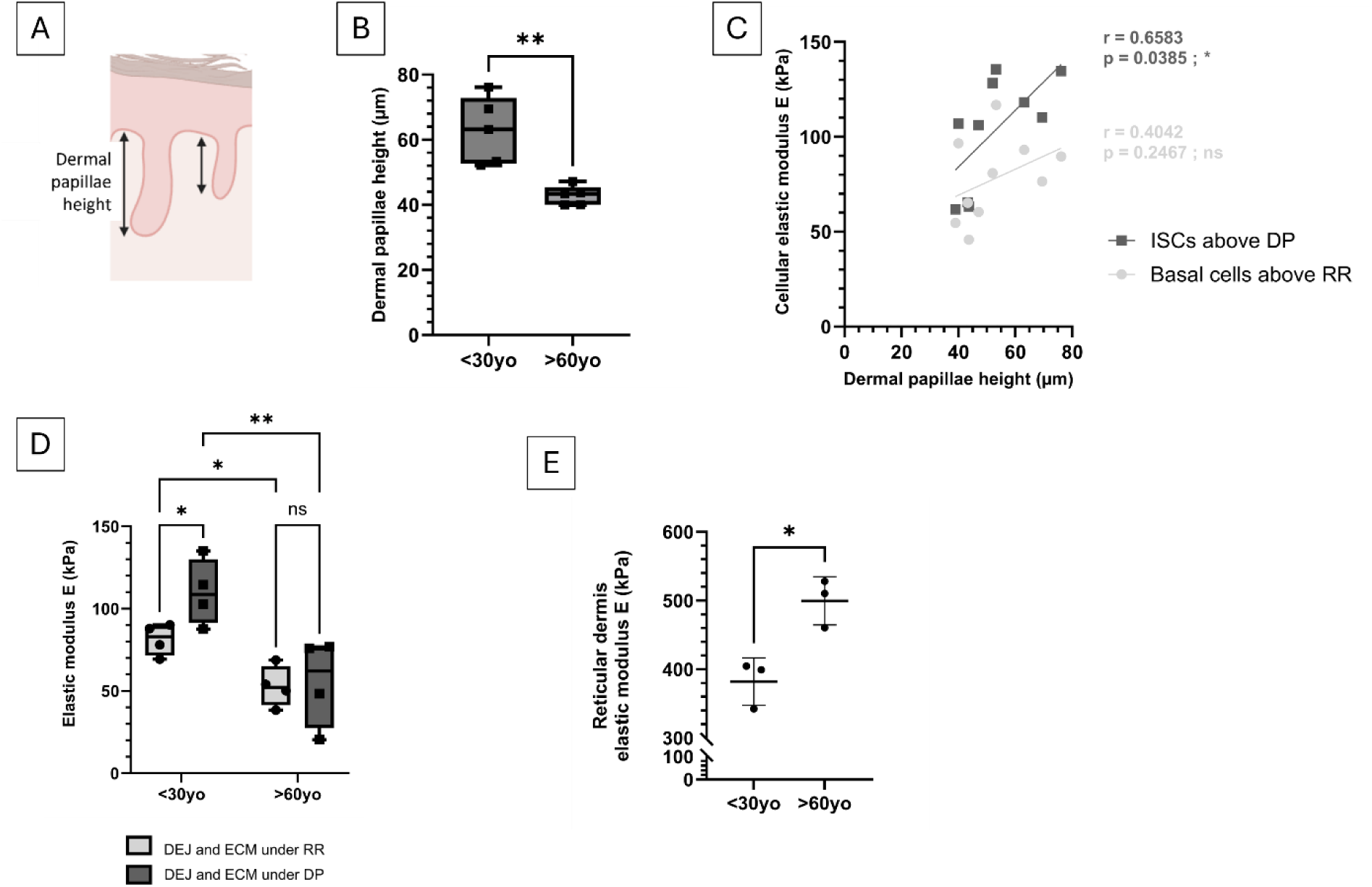
**(A)** Schematic representation of the method used to analyze DP height. **(B)** Quantification of DP height in human skin sections from donors under 30 years old (N=5) compared to those over 60 years old (N=5). An unpaired t-test was performed to compare DP height between the two age groups (** p-value < 0.005). **(C)** Elastic modulus of ISCs and basal cells above the RR in relation to DP height for donors under 30 years (N=5) and over 60 years (N=5). A Pearson correlation analysis was conducted to assess the relationship between elastic modulus of basal cells and changes in DP height (* p-value < 0.05). **(D)** Box plot depicting the average stiffness of the ECM under ISCs above the DP (dark grey) and the ECM under RR (light grey), categorized by age. The younger group included donors under 30 years (N=4; ages: 22, 25, 27, 29), while the older group comprised donors over 60 years (N=4; ages: 60, 61, 62, 64). Each donor provided three sections, with over 15 AFM measurements taken for each section. A two-way ANOVA with Tukey’s multiple comparison test was employed to compare stiffness between ISCs and basal cells above the RR across age groups (ns = not significant; * p-value < 0.05; ** p-value < 0.01). **(E)** Plot illustrating the average stiffness of the reticular dermis according to age, comparing young donors under 30 years (N=3) to older donors over 60 years (N=3). Each donor contributed three sections, with more than 20 AFM measurements donors. A t-test was performed to assess differences (* p-value < 0.05).

### Alterations in the mechanical environment of ISCs during aging

The DEJ and ECM proteins beneath it serve as substrates for ISCs within their environment. Research indicates that substrate stiffness influences cell fate and the mechanical properties of overlying cells [44], [45]. To explore this, we conducted AFM measurements on the DP or RR regions, specifically examining the subjacent ECM proteins beneath ISCs expressing MCSP and beneath basal cells lacking MCSP in cryosections of young human skin. In skin samples from individuals under 30 years of age (Figure 2D), we observed a notable difference in the stiffness of DEJ and ECM proteins based on location. The average stiffness of DP and ECM proteins beneath ISCs was significantly higher than that of RR and ECM proteins beneath basal cells. To assess the impact of aging on ECM stiffness, we performed similar AFM measurements on cryosections from individuals over 60 years old (Figure 2D). In this group, no significant differences in the stiffness of DEJ and ECM proteins were found based on location; the average stiffness beneath ISCs in the DP was comparable to that observed beneath basal cells in the RR. Furthermore, we noted a significant decrease in the stiffness of DP and ECM proteins under ISCs with age. Conversely, the rigidity of the reticular dermis increased with aging (Figure 2E). These findings indicate that a specific mechanical environment exists beneath ISCs in skin from individuals under 30 years old, whereas this specific mechanical environment is diminished in older skin.

### MCSP as cell sorting marker for isolating ISCs located above the DP

To investigate the mechanical properties of ISCs located above the DP *in vitro*, we developed a sorting technique utilizing the stem cell marker MCSP. Freshly isolated keratinocytes from human skin biopsies of varying ages were sorted based on the expression of markers CD49, CD71, and MCSP. This approach allowed us to categorize different basal cell populations: basal CD49high-CD71low-MCSPlow stem cells designated as MCSP-SCs; transient CD49high-CD71high-MCSPhigh amplifying cells referred to as MCSP+ TA; and CD49high-CD71low-MCSPhigh ISCs above the DP termed MCSP+ SCs. Initially, after selecting viable cells organized into singlets, we found that only 18.5% of the SCs from a young donor exhibited high levels of MCSP (Figure 3A). Quantifying data from all young donors revealed an average of approximately 13% MCSP+ SCs (Figure 3B). In contrast, the average frequency of MCSP+ SCs in the elderly group was lower, although this difference was not statistically significant (Figure 3B). In the young cohort, analysis of cell size using the MFI FSC-A parameter revealed that MCSP+ SCs seems to exhibit a higher FSC-A than MCSP-SCs (Figure 3C). This size distinction was not observed in the older group (data not shown), suggesting that ISCs above DP are larger than other epidermal stem cells outside the DP apex, with this size advantage diminishing with age. Regarding MCSP + TA cells, a significant size difference was noted between MCSP+ SCs and MCSP + TA cells in the young donors (Figure 3D). However, this size disparity was not present in the elderly group, although a similar trend was observed (data not shown). Subsequently, the sorted cells were cultured on a rat collagen I matrix at 100µg/ml. After one hour of culture, we assessed cell size, revealing that the area of MCSP+ SCs tend to be larger than that of MCSP-SC, and smaller than that of MCSP + TA cells (Figure 3E). After one and two days in culture, we noted that, in one donor of 22 years, MCSP+ SCs appeared to spread more rapidly than MCSP-SCs, which remained relatively round in the unsorted cells (Figure 3F). These observations suggest that MCSP+ SCs seems to exhibit distinct size and spreading capabilities compared to MCSP-SCs.

**Figure 3.**
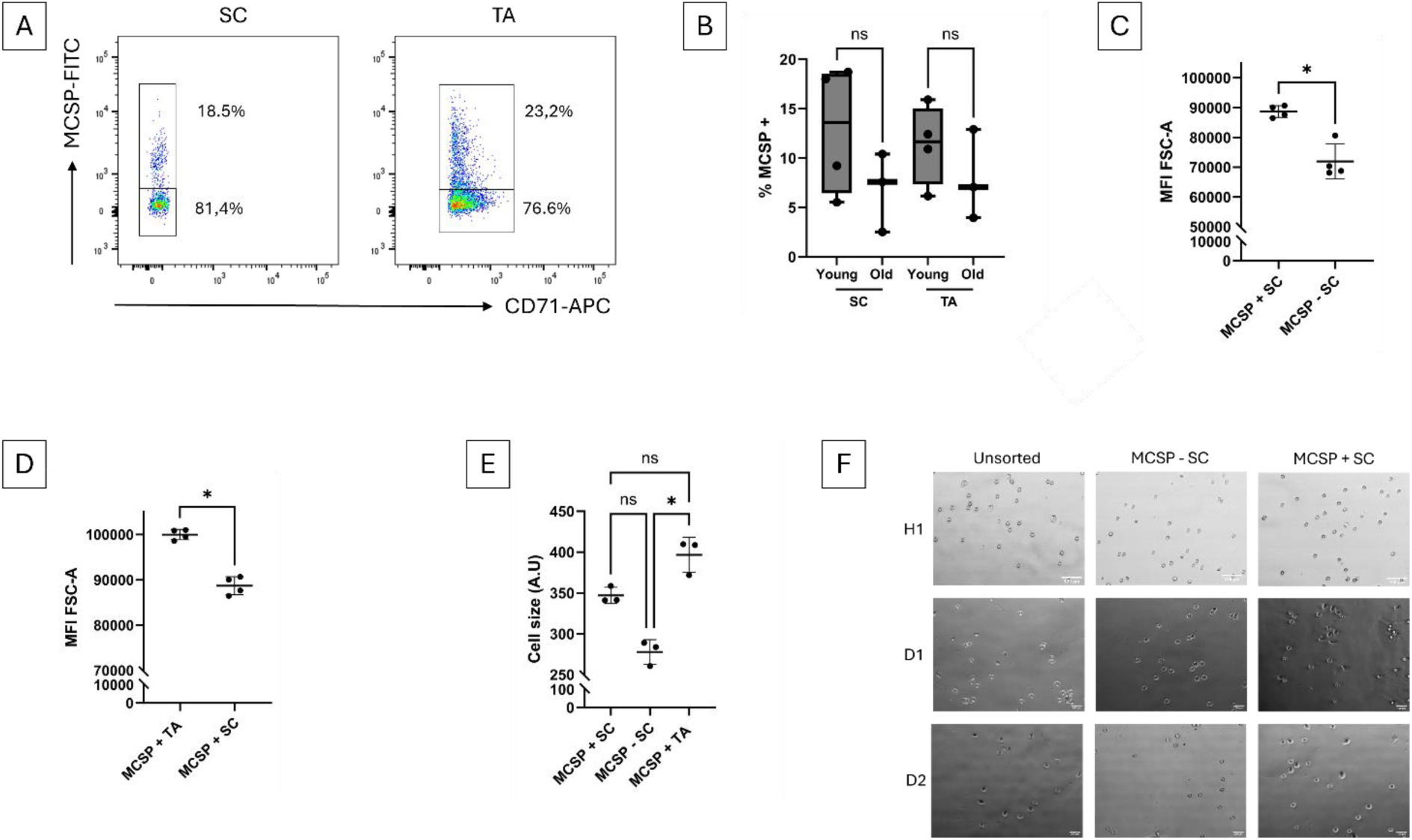
A, B, C and D, flow cytometric analysis of CD49, CD71, and MCSP protein expression in primary human keratinocytes (P0) isolated from human skin of varying ages. The young skin group included donors aged 20, 24, 31, and 34 years (N=4), while the old skin group comprised donors aged 62, 70, and 78 years (N=3). Staining and acquisitions were from a single experiment. A Mann-Whitney test was used for comparisons across conditions (* p-value < 0.05). **E** and **F**, SC and TA cells were FACS sorted based on their expression of MSCP and cultured on a rat collagen I matrix for 1 hour, 1 day and 2 days. **(A)** Flow cytometric plots showing MSCP expression in stem cells (SC: CD49high CD71low cells) and TA cells (TA: CD49high CD71high cells) from a representative young donor. **(B)** Box plot illustrating the proportions of MCSP positive cells among SC and TA cells for both young and old donors. **(C)** Plot showing the comparison of mean FSC-A of SC MCSP + SC *vs* MCSP + TA in the young donors. **(D)** Plot showing the comparison of mean FSC-A of MCSP + SC *vs* MCSP + SC in the young donors. **(E)** Cell size analysis after 1 hour culture on a rat collagen I matrix (100µg/ml) for sorted cells isolated from human skin (N=3; ages: 22, 37, and 44 years). **(F)** Morphological images of a 22-year-old donor illustrating unsorted cells, MCSP - SC and MCSP + SC on a rat collagen I matrix (100µg/ml) after 1 hour, 1 day, and 2 days of culture.

## DISCUSSION

Epidermal renewal and skin homeostasis are maintained by basal layer ISCs situated on the DEJ [1]. During aging, a decline in epidermal renewal correlates with a reduction in the stem potential of ISCs and alterations in the protein composition of the DEJ and dermis, contributing to the skin changes observed with age [46] [35]. The study of cellular mechanical properties can serve as a means to identify cell types [47], [48]. Furthermore, cell fate and cell’s mechanical property can be directly influenced by the rigidity of its substrate [44], [45]. In this study, we propose employing AFM to characterize the mechanical signature of ISCs during aging, as well as those of their immediate environment, the DEJ.

To locate putative ISCs in the basal layer, we first conducted an immunofluorescence-based approach on all human skin sections, utilizing MCSP labeling, which is recognized as a stem cell marker. This approach confirmed the expected localization of ISCs in the basal layer above the DPs. According to existing literature, ISCs are specifically positioned above the DP and express the stem cell marker β1-integrin, plays a critical role in regulating the balance between stem cell renewal and differentiation [20]. Basal layer cells that express high levels of β1-Integrin also express the MCSP protein [9], [33]. The precise localization of MCSP enabled us to accurately identify ISCs above the DP in all donor samples.

Using AFM, we demonstrated that putative ISCs located above the DP and expressing MCSP possess a distinct mechanical signature in young skin. Specifically, these ISCs exhibit greater cell stiffness compared to basal layer cells situated above the RR. The stiffness measurements we obtained account for various factors, including cells junctions and cytoskeletal organization. It is well established that stem cells are organized within a niche that provides a specific microenvironment [49]. ISCs are characterized by a high density of cell-cell junctions and robust cell-basement membrane attachments to ECM proteins such as laminin-332, facilitated through integrins [20]. These interactions play a critical role in maintaining the stem cell niche and regulating the differentiation of epidermal ISCs [17], [21]. Furthermore, proteins like α6-integrin, type XVII collagen, and laminin-332, contribute to hemidesmosomes [50], [51] that anchor ISCs to the DEJ, influencing the niche’s integrity and behavior via mechano-transduction pathways, such as YAP-TAZ signaling [52]. The junctions present in ISCs are linked to their cytoskeleton, particularly through integrins, which connect to and remodel the actin cytoskeleton [39], [53]. Cytoskeletal organization significantly influences cell stiffness [39], [54], [55] However, our AFM observations in aged skin reveal that ISCs lose their distinct mechanical signature, indicating that aging adversely affects the mechanical properties of ISCs. In older individuals, ISCs exhibit stiffness comparable to that of basal cells above the RR, with an overall decrease in the stiffness of all basal epidermal cells observed during aging. This loss of mechanical distinctiveness in ISCs may be attributed to changes in cell fate and microenvironment. As individuals age, ISCs show a decline in their renewal potential, which is accompanied by reductions in stem cell markers such as α6, β1-integrin, and MCSP [8], [32], [33]. Additionally, aging is associated with decreased expression of ECM proteins like laminin-332 and collagen VII, leading to a reduction in hemidesmosome size between ISCs and the DEJ [31], [56]. We propose that these modifications adversely impact the cytoskeletal organization of ISCs, resulting in diminished cellular rigidity.

The observed differences in stiffness properties of ISCs, consistently located above the DP, can be attributed to the distinct topography of the DP compared to the RR. As anticipated, we noted a decrease in DP height in aged skin, which correlates with the flattening of the DEJ [32]. Our findings reveal a positive correlation between DP height and ISCs stiffness specifically, as DP height increases, stiffness of ISCs also rises. An *in vitro* study has shown that substrate topography influences the behavior of epidermal ISCs, potentially promoting or inhibiting their differentiation by regulating the organization of the actin cytoskeleton [43]. Furthermore, research indicates that the patterning of epidermal stem cells is influenced by mechanical forces exerted at intercellular junctions in response to the undulations of the DEJ [57]. Additionally, Viswanathan *et al.* demonstrated that stem cells cultured on substrates with wavy topography express either stem cell markers or differentiation markers based on their positioning at the peaks of the waves [58].

Given that keratinocytes are mechanosensitive cells capable of detecting the mechanical properties of their environment [59], [60], we also focused on the mechanical characteristics surrounding ISCs. These cells rest on the DEJ and are indirectly supported by the papillary dermis, prompting us to investigate their stiffness using AFM. Our findings reveal that the protein components of the DEJ and a portion of the papillary dermis beneath the putative ISCs exhibit greater stiffness compared to those located beneath basal layer cells above the RR. Existing literature indicates that substrate rigidity significantly influences cell rigidity, primarily by affecting cytoskeletal organization [45]. Furthermore, substrate stiffness is directly correlated with the proliferation capacity of keratinocytes, which tends to increase on more rigid substrates [24], [25]. In this context, we have identified a unique mechanical signature of ISCs that may stem from the high stiffness of their underlying substrate. Aging, however, is associated with a loss of this specific mechanical signature in the ECM beneath ISCs. This change may be attributed to a reduction in ECM protein component, such as laminin-332 and collagen VII [31], [56], as well as a decrease in the diameter of collagen bundles [61]. Each ECM protein possesses its own stiffness [62], which can influence the overall stiffness of the ECM. We propose that alterations in DEJ and ECM components during aging affect the rigidity of the substrate within ISCs niches, consequently modulating their cellular stiffness. Moreover, our findings corroborate previous reports of increased stiffness in the reticular dermis with age [63]. Thus, we postulate that the observed decrease in stiffness at the DEJ beneath ISCs during aging reflects a dynamic mechanical environment that evolves in conjunction with the mechanics of the ISCs themselves.

Although AFM provides valuable insights into the biomechanics of epidermal basal cells in human skin sections, it is essential to isolate stem cells to better characterize their properties *in vitro*. Therefore, our objective was to sort stem cells based on their expression of CD49, CD71 [13], [14], [15], and MCSP to specifically isolate the stem cells located above the DP that express high levels of MCSP. Our findings reveal that around only 12% of the stem cells express high level of MCSP in young donors. This result can be attributed to the considerable heterogeneity present within the basal layer stem cell population at various transitional states [12]. Additionally, the non-significant decrease in the average number of MCSP+ SCs observed during aging may be explained by a decline in MCSP expression [33]. In young donors, we noted that the MCSP+ SCs situated above the DP seems larger than the MCSP-SCs but smaller than TA cells. Cell size can be indicative of cell fate; for instance, quiescent epidermal SCs are typically smaller than proliferative TA cells [15]. Therefore, we hypothesize that MCSP+ SCs do not exhibit a transient amplifying cell profile. Given the presence of distinct stem cell populations at various transition stages [12] and the larger size of MCSP+ SCs compared to MCSP-SCs, we suggest that MCSP+ SCs represent a specific subpopulation of basal layer stem cells. Interestingly, we observed that in one donor, MCSP+ SCs spread more rapidly than both MCSP-SCs and unsorted cells on a collagen matrix. This may be attributed to the role of MCSP in modulating cell adhesion [16], [64], supporting the notion that MCSP+ SCs possess unique characteristics as a subpopulation of basal layer stem cells. However, further characterization of isolated MCSP+ SCs is necessary, including the examination of their expression of additional stem cell markers in culture, their ability to form clones, and an assessment of their cellular rigidity *in vitro*.

Sorting cells expressing CD49high, CD71low, and MCSPhigh enables the specific isolation of a subpopulation of epidermal stem cells, namely MCSP-positive stem cells located above the DP. This targeted approach facilitates a more detailed mechanical characterization of these cells and aids in understanding the mechanical deficiencies associated with impaired epidermal renewal during aging. By AFM we revealed that ISCs and the DEJ possess a specific mechanical signature lost during aging. Employing AFM as a tool to assess the mechanical signature of ISCs can extend to the evaluation of reconstructed skin enriched with stem cells, as well as skin exhibiting renewal challenges, such as diabetic and psoriatic conditions. This application opens new avenues for utilizing AFM-based techniques as complementary tools for monitoring therapies designed to enhance skin cell renewal and overall skin health.

## Abbreviations

DEJ: Dermo-Epidermal Junction

ECM: Extra Cellular Matrix

RR: Rete Ridge

DP: Dermal Papilla

ISC: Interfollicular Stem Cells

TA: Transit Amplifying cells

YO: Years Old

## Acknowledgments

We acknowledge the contribution of SFR Biosciences (Université Claude Bernard Lyon 1, CNRS UAR3444, Inserm US8, ENS de Lyon) for the help of the staff of - AniRA Cytometry core facility, especially Estelle Devêvre and Tiffany Deborde, for assistance with Fluorescence Activated Cell Sorting (FACS).

## Author contributions

C.B., S.M., JE.L., J.C. and G.R. designed research; S.M. performed experiments; C.B., S.M., JE.L., J.C. and G.R., contributed to analysis and interpretation of data; S.M. and C.B. wrote the paper; G.R. edited the manuscript.

---

**Figure supp 1.**
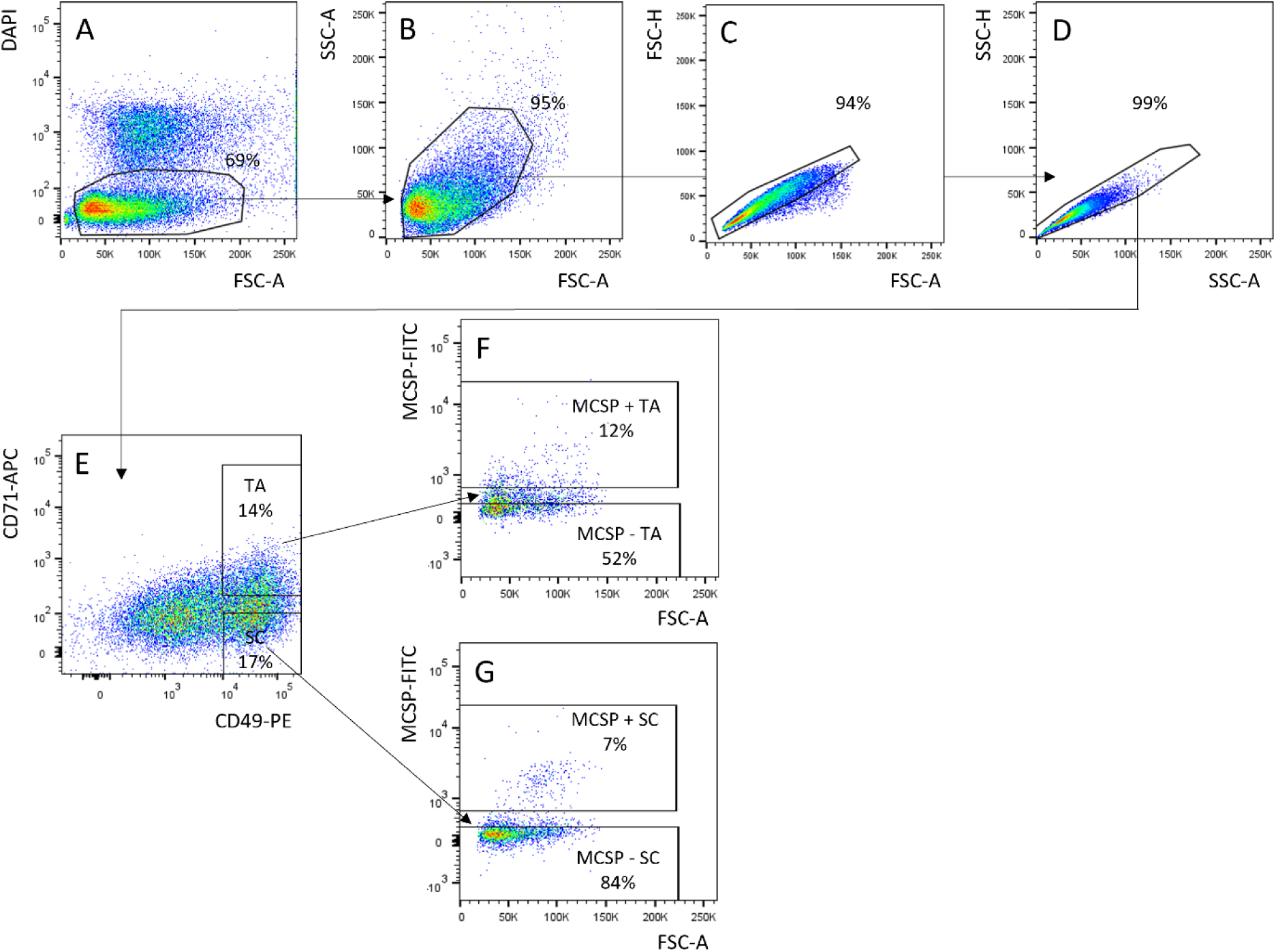
Gating strategy for Fluorescence Activated Cell Sorting (FACS) of P0 primary human keratinocytes isolated from human skins of different ages. First, live cells are selected based on their non incorporation of DAPI (**A**), then morphological intact cells are gated based on the physical parameters Side Scatter-Area (SSC-A) and Forward Scatter-A (FSC-A) (**B**) and single cells are identified by comparing FSC-Height vs FSC-Area followed by SSC-Height vs SSC-Area (**C and D**). Finally, among live single cells, SC (CD49high CD71low) and TA (CD49high CD71high) are identified (**E**) and each population is divided into 2 based on their level of expression of MCSP for 4 sorting subsets: MCSP + SC and MCSP - SC (**F**) and MCSP + TA and MCSP - TA (**G**).

